## Supplementary Material for "Loss of SET1/COMPASS methyltransferase activity reduces lifespan and fertility in *Caenorhabditis elegans*"

### Supplementary Information

#### Immunoprecipitation for proteomics

Immunoprecipitations were performed on frozen embryos prepared by hypochlorite treatment from animals grown at 20°C on enriched NGM seeded with concentrated HB101 bacteria. After hypochlorite treatment embryos were washed once in IP buffer (50 mM HEPES/KOH pH 7,5; 300 mM KCl; 1 mM EDTA; 1 mM MgCl2; 0.2% Igepal-CA630 and 10% glycerol) and flash-frozen in beads in liquid nitrogen. Embryos were then grounded to powder, resupended in IP buffer containing complete protease inhibitors (Roche) and sonicated on ice to an amplitude of 30% for 2.5 min (15'' ON/ 15''OFF pulses) using an Ultrasonic Processor (Bioblock Scientific). Protein extracts were recovered in the supernantant after centrifugation at 20,000 g for 15 min at 4°C. Protein concentration was estimated using the Bradford assay (Bio-Rad Protein Assay Dye).

For all GFP immunoprecipitations, 70 mg of total protein extract was incubated for preclearing with 200µL slurry of binding control magnetic agarose beads (Chromotek bmab-20) in IP buffer for 1h at 4°C. Then 200 µl of GFP-TRAP MA beads slurry (Chromotek) were added to each sample and incubation continued for additional 3h on a rotator. Beads were collected with a magnet, washed three times in IP buffer and once in Benzo buffer (HEPES/KOH 50 mM pH 7,5; KCl 150 mM; EDTA 1 mM; $MgCl_2$ 1 mM; Igepal-CA630 0.2%; glycerol 10%). Beads were then incubated in 400 µl of Benzo buffer containing 2,500 units of benzonase (Sigma) for 1 h at 4°C and washed three times in IP buffer. Eluates were recovered by incubation at 95°C for 10 min in 60 µl of 1x LDS buffer (Thermo Fischer). 1/10 of each eluate was resolved on a 4–12% NuPage Novex gel (Thermo Fischer) and stained with SilverQuest staining kit (Thermo Fischer). 40 µl of the eluates was then analyzed by mass spectrometry.

For HA::WDR-5.1 co-immunoprecipitation, samples were prepared as described previously (1).

**Mass spectrometry-based proteomic analyses**

For the GFP co-immunoprécipitation: The eluted proteins were stacked at the top of a 4-12% NuPAGE gel (Invitrogen). After staining with R-250 Coomassie Blue (Biorad), proteins were digested in-gel using modified trypsin (sequencing purity, Promega), as previously described (2). The resulting peptides were analyzed by online nanoliquid chromatography coupled to MS/MS (Ultimate 3000 RSLCnano and Q-Exactive HF, Thermo Fisher Scientific) using a 120 min gradient. To this end, peptides were sampled on a precolumn (300 µm x 5 mm PepMap C18, Thermo Scientific) and separated in a 75 µm x 250 mm C18 column (Reprosil-Pur 120 C18-AQ, 1.9 µm, Dr. Maisch). The MS and MS/MS data were acquired by Xcalibur (Thermo Fisher Scientific).

Peptides and proteins were identified by Mascot (version 2.7.0, Matrix Science) through concomitant searches against the Uniprot database (*C. elegans* taxonomy, January 2021 version), classical contaminants database (homemade), and the corresponding reversed databases. Trypsin/P was used for digestion and two missed cleavages were allowed. Precursor and fragment mass error tolerances were set at 10 and 20 ppm, respectively. Peptide modifications allowed during the search were: Carbamidomethyl (C, fixed), Acetyl (Protein N-term, variable) and Oxidation (M, variable). The Proline software (3) was used for the compilation, grouping and filtering of the results (conservation of rank 1 peptides, peptide length ≥ 6 amino acids, peptide score ≥ 25, and false discovery rate of peptide-spectrum-match identifications < 1% as calculated on peptide-spectrum-match scores by employing the reverse database strategy). Proline was then used to perform a compilation, grouping and spectral counting-based comparison of the protein groups identified in the different samples. Proteins from the contaminant database were discarded from the final list of identified proteins. To be considered as a potential CFP-1 interactor, a protein must be identified with a minimum of 3 specific spectral counts and not in the negative control eluate, or enriched at least 5 times compared to the negative control eluate.

For the HA::WDR-5.1 mass spectrometry, samples were prepared and analysed as described previously (1).

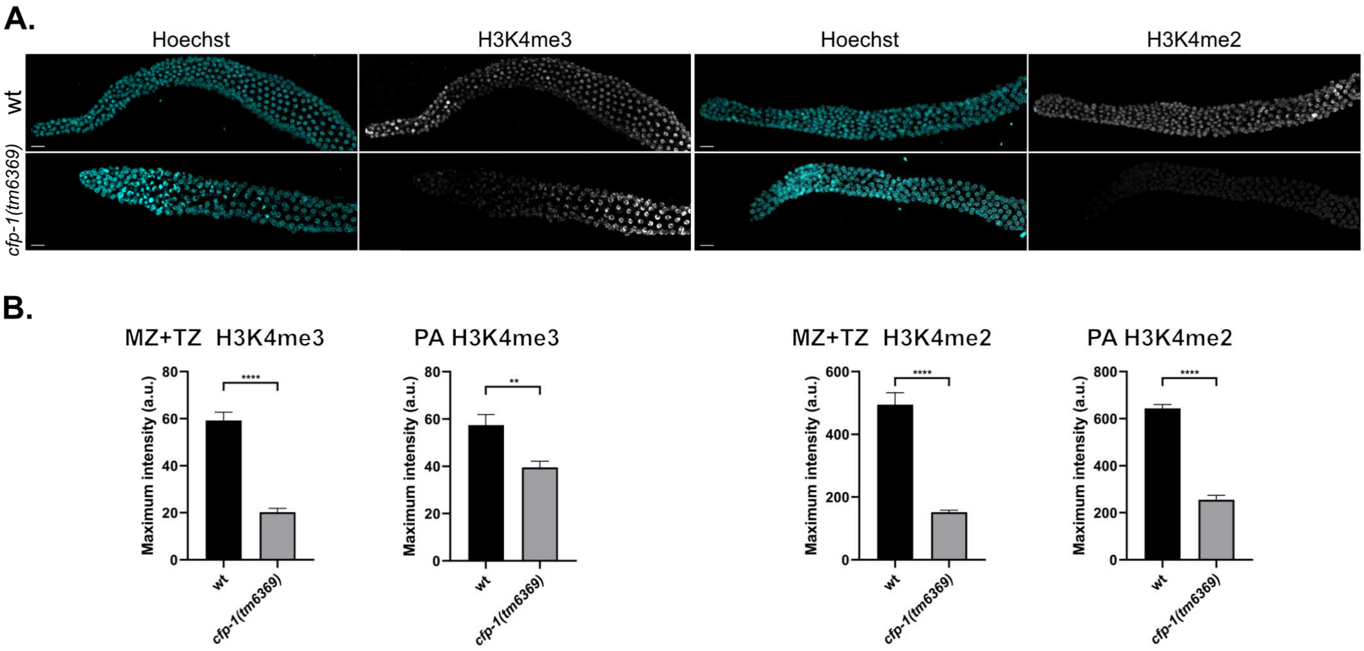


**Figure S1.** Confocal images of dissected gonads from wild type and *cfp-1(tm6369)* mutants stained with Hoechst and anti-H3K4me3 antibody (left panels), or Hoechst and anti-H3K4me2 (right panels). B. Quantifications of the H3K4me3/2 signals in MZ+TZ and PA were performed using mean pixel intensities; error bars show SEM, n=5 (P<0.001, P**<0.0001, mutant vs wild type worms, unpaired t-test).

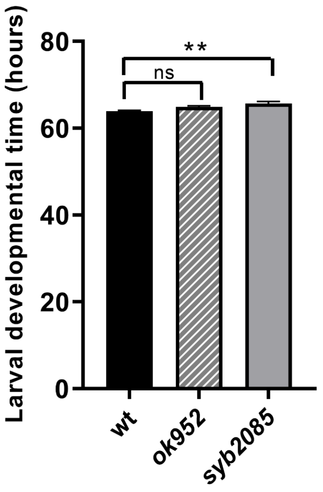


**Figure S2.** Larval developmental time (egg to adult) measured for *set-2(ok952)* and *set-2(syb2085)* mutants, and wild type; error bars show SEM, n=10 (P**<0.001, *set-2(syb2085)* vs wild type worms, one-way ANOVA folllowed by Dunnett's test).

**Table S1.** List of SET-2 peptides identified by mass spectrometry in the GFP::CFP-1 immunoprecipitates obtained from embryos carrying the indicated *set-2* alleles.

| Co-IP GFP::CFP-1 | | | | | | | | | | | |
|---|---|---|---|---|---|---|---|---|---|---|---|
| *set-2(+)* | | | | *set-2 (ok952)* | | | | *set-2 (syb2085)* | | | |
| SET-2 Peptides | Start | Stop | Score | SET-2 Peptides | Start | Stop | Score | SET-2 Peptides | Start | Stop | Score |
| | | | | | | | | WAWQKVFETGK | 37 | 47 | 39,24 |
| DGFPQDCKSKEDFER | 53 | 67 | 66,8 | DGFPQDCKSKEDFER | 53 | 67 | 60,44 | DGFPQDCKSKEDFER | 53 | 67 | 71,22 |
| | | | | | | | | KNFESLQQSSVYQTNSFR | 86 | 103 | 49,73 |
| | | | | KNFESLQQSSVYQTNSFRNPR | 86 | 106 | 31,85 | KNFESLQQSSVYQTNSFRNPR | 86 | 106 | 34,57 |
| NFESLQQSSVYQTNSFR | 87 | 103 | 71,67 | NFESLQQSSVYQTNSFR | 87 | 103 | 35,26 | NFESLQQSSVYQTNSFR | 87 | 103 | 119,12 |
| NFESLQQSSVYQTNSFRNPR | 87 | 106 | 67,6 | NFESLQQSSVYQTNSFRNPR | 87 | 106 | 30,14 | NFESLQQSSVYQTNSFRNPR | 87 | 106 | 55,58 |
| VDSYYCTIPPKR | 115 | 126 | 59,66 | VDSYYCTIPPKR | 115 | 126 | 59,36 | VDSYYCTIPPKR | 115 | 126 | 65,5 |
| EVSLFNMDDNCTEVLLR | 127 | 143 | 75,79 | EVSLFNMDDNCTEVLLR | 127 | 143 | 53,87 | EVSLFNMDDNCTEVLLR | 127 | 143 | 96,96 |
| EAHNFYSMYHAQNLLATK | 179 | 196 | 35,31 | EAHNFYSMYHAQNLLATK | 179 | 196 | 30,46 | EAHNFYSMYHAQNLLATK | 179 | 196 | 43,79 |
| EAHNFYSMYHAQNLLATK | 179 | 196 | 28,06 | ANFLRDQNEKYELAMR | 241 | 256 | 29,87 | EAHNFYSMYHAQNLLATK | 179 | 196 | 29,11 |
| | | | | ANFLRDQNEKYELAMR | 241 | 256 | 27,68 | ANFLRDQNEKYELAMR | 241 | 256 | 77,12 |
| | | | | DQNEKYELAMR | 246 | 256 | 36,45 | ANFLRDQNEKYELAMR | 241 | 256 | 52,05 |
| NANGDVVKYETYKMEK | 407 | 422 | 28,78 | NANGDVVKYETYK | 407 | 419 | 44,8 | NANGDVVKYETYK | 407 | 419 | 37,45 |
| | | | | | | | | NANGDVVKYETYKMEK | 407 | 422 | 25,68 |
| | | | | | | | | GKNQLENVSSESASGSSSVDTYPDFSDEER | 448 | 477 | 56,96 |
| ELSSTHTNSVPNLK | 536 | 549 | 52,41 | ELSSTHTNSVPNLK | 536 | 549 | 57,23 | ELSSTHTNSVPNLK | 536 | 549 | 60,99 |
| FSGIFGPTQR | 684 | 693 | 69,06 | FSGIFGPTQR | 684 | 693 | 46,98 | FSGIFGPTQR | 684 | 693 | 44,41 |
| EEPAQVEVEYDYPLK | 694 | 708 | 95,67 | | | | | FSGIFGPTQREEPAQVEVEYDYPLK | 684 | 708 | 38,12 |
| | | | | | | | | EEPAQVEVEYDYPLK | 694 | 708 | 27,77 |
| KVAEDIRQQIMR | 775 | 786 | 42,22 | KVAEDIRQQIMR | 775 | 786 | 42,75 | KVAEDIRQQIMR | 775 | 786 | 42,25 |
| KVAEDIRQQIMR | 775 | 786 | 29,64 | KVAEDIRQQIMR | 775 | 786 | 28,43 | KVAEDIRQQIMR | 775 | 786 | 36,61 |
| QCFAALDEKLHLK | 787 | 799 | 25,89 | QCFAALDEKLHLK | 787 | 799 | 43,2 | QCFAALDEK | 787 | 795 | 39,05 |
| | | | | | | | | QCFAALDEKLHLK | 787 | 799 | 36,49 |
| ARQEAEKPSNHLIADMMTLYNNQSFASSSR | 815 | 844 | 45,56 | | | | | ARQEAEKPSNHLIADMMTLYNNQSFASSSR | 815 | 844 | 29,88 |
| QEAEKPSNHLIADMMTLYNNQSFASSSR | 817 | 844 | 46,25 | | | | | QEAEKPSNHLIADMMTLYNNQSFASSSR | 817 | 844 | 48,18 |
| | | | | | | | | QEAEKPSNHLIADMMTLYNNQSFASSSR | 817 | 844 | 27,86 |
| ASFSSTSIQSSPER | 940 | 953 | 78,54 | | | | | ASFSSTSIQSSPER | 940 | 953 | 102,65 |
| KLIMSSDESSTTGSTATSVVSSR | 986 | 1008 | 89,82 | | | | | RKLIMSSDESSTTGSTATSVVSSR | 985 | 1008 | 49,97 |
| LIMSSDESSTTGSTATSVVSSR | 987 | 1008 | 207,5 | **deletion in *set-2 (ok952)*** | | | | KLIMSSDESSTTGSTATSVVSSR | 986 | 1008 | 86,37 |
| LIMSSDESSTTGSTATSVVSSR | 987 | 1008 | 185,01 | **No peptide found in MS** | | | | KLIMSSDESSTTGSTATSVVSSR | 986 | 1008 | 60,59 |
| | | | | | | | | LIMSSDESSTTGSTATSVVSSR | 987 | 1008 | 227,79 |
| | | | | | | | | LIMSSDESSTTGSTATSVVSSR | 987 | 1008 | 94,49 |
| SQTDFISER | 1028 | 1036 | 45,24 | | | | | SQTDFISER | 1028 | 1036 | 59,27 |
| SQTDFISERVSK | 1028 | 1039 | 25,02 | | | | | SQTDFISERVSK | 1028 | 1039 | 46,58 |
| IEGEERPLPEPVETSGPIIGDSSYLPYK | 1040 | 1067 | 59,28 | | | | | IEGEERPLPEPVETSGPIIGDSSYLPYK | 1040 | 1067 | 124,13 |
| AGIIEMNLPANSIR | 1074 | 1087 | 73,88 | AGIIEMNLPANSIR | 1074 | 1087 | 62,29 | AGIIEMNLPANSIR | 1074 | 1087 | 83,6 |
| AGIIEMNLPANSIR | 1074 | 1087 | 53,43 | AGIIEMNLPANSIR | 1074 | 1087 | 53,45 | AGIIEMNLPANSIR | 1074 | 1087 | 80,5 |
| | | | | AHEYHPFTTEHCYFGIDDPRQPK | 1088 | 1110 | 39,59 | AHEYHPFTTEHCYFGIDDPRQPK | 1088 | 1110 | 44,28 |
| | | | | | | | | IQIFDHSPCKSEPGSEPLK | 1111 | 1129 | 57,26 |
| IQIFDHSPCK | 1111 | 1120 | 43,95 | IQIFDHSPCK | 1111 | 1120 | 41,5 | IQIFDHSPCK | 1111 | 1120 | 57,23 |
| KQVFEKDPYEYYEPPPTK | 1167 | 1184 | 28,31 | KQVFEKDPYEYYEPPPTK | 1167 | 1184 | 75,45 | KQVFEKDPYEYYEPPPTK | 1167 | 1184 | 105,03 |
| | | | | | | | | KIIGDCEDLPDLEDQWYLR | 1206 | 1224 | 28,23 |
| | | | | | | | | AALNEMQSEVKSADELPWKK | 1225 | 1244 | 75,05 |
| AALNEMQSEVK | 1225 | 1235 | 66,32 | AALNEMQSEVK | 1225 | 1235 | 45,77 | AALNEMQSEVK | 1225 | 1235 | 43,77 |
| AALNEMQSEVKSADELPWKK | 1225 | 1244 | 25,35 | AALNEMQSEVKSADELPWKK | 1225 | 1244 | 41,46 | AALNEMQSEVKSADELPWK | 1225 | 1243 | 30,12 |
| | | | | SADELPWK | 1236 | 1243 | 44,96 | AALNEMQSEVKSADELPWKK | 1225 | 1244 | 25,76 |
| | | | | KMLTFKEMLR | 1244 | 1253 | 27,46 | SADELPWK | 1236 | 1243 | 45,81 |
| MLTFKEMLR | 1245 | 1253 | 51,59 | MLTFKEMLR | 1245 | 1253 | 55,2 | MLTFKEMLR | 1245 | 1253 | 52,24 |
| MLTFKEMLR | 1245 | 1253 | 35,47 | MLTFKEMLR | 1245 | 1253 | 40,78 | MLTFKEMLR | 1245 | 1253 | 51,05 |
| MLTFKEMLR | 1245 | 1253 | 29 | MLTFKEMLR | 1245 | 1253 | 35,58 | MLTFKEMLR | 1245 | 1253 | 44,81 |
| SEDPLLRLNPIR | 1254 | 1265 | 44,53 | SEDPLLRLNPIR | 1254 | 1265 | 46,03 | MLTFKEMLR | 1245 | 1253 | 35,74 |
| KGLPDAFYEDEELDGVIPVAAGCSR | 1268 | 1292 | 94,64 | KGLPDAFYEDEELDGVIPVAAGCSR | 1268 | 1292 | 122,7 | SEDPLLRLNPIR | 1254 | 1265 | 44,57 |
| GLPDAFYEDEELDGVIPVAAGCSR | 1269 | 1292 | 39,15 | GLPDAFYEDEELDGVIPVAAGCSR | 1269 | 1292 | 25,67 | KGLPDAFYEDEELDGVIPVAAGCSR | 1268 | 1292 | 144,15 |
| RPDNESHPTAIFSERDETAIR | 1310 | 1330 | 27,72 | RPDNESHPTAIFSERDETAIR | 1310 | 1330 | 69,6 | RPDNESHPTAIFSERDETAIR | 1310 | 1330 | 54,67 |
| RPDNESHPTAIFSER | 1310 | 1324 | 64,89 | RPDNESHPTAIFSER | 1310 | 1324 | 40,08 | RPDNESHPTAIFSER | 1310 | 1324 | 37,09 |
| RLLTSLGDANNDFFK | 1345 | 1359 | 44,51 | LLTSLGDANNDFFK | 1346 | 1359 | 114,62 | RLLTSLGDANNDFFKINQLK | 1345 | 1364 | 46,78 |
| GIGSSYLFR | 1418 | 1426 | 64,53 | RGIGSSYLFR | 1417 | 1426 | 28,22 | RLLTSLGDANNDFFK | 1345 | 1359 | 45,09 |
| | | | | GIGSSYLFR | 1418 | 1426 | 74,02 | LLTSLGDANNDFFK | 1346 | 1359 | 114,26 |
| | | | | IDLHHVIDATKR | 1427 | 1438 | 45,8 | RGIGSSYLFR | 1417 | 1426 | 34,57 |
| IDLHHVIDATK | 1427 | 1437 | 47,56 | IDLHHVIDATK | 1427 | 1437 | 41,06 | GIGSSYLFR | 1418 | 1426 | 73,93 |
| | | | | FINHSCQPNCYAK | 1444 | 1456 | 28,78 | IDLHHVIDATK | 1427 | 1437 | 61,11 |
| | | | | VLTIEGEKRIVIYSR | 1457 | 1471 | 29,41 | IDLHHVIDATKR | 1427 | 1438 | 27,5 |
| TIIKKGEEITYDYK | 1472 | 1485 | 55,9 | TIIKKGEEITYDYK | 1472 | 1485 | 70,51 | TIIKKGEEITYDYK | 1472 | 1485 | 51,44 |
| GEEITYDYK | 1477 | 1485 | 40,49 | GEEITYDYK | 1477 | 1485 | 35,9 | GEEITYDYK | 1477 | 1485 | 31,54 |
| FPIEDDKIDCLCGAK | 1486 | 1500 | 46,77 | FPIEDDKIDCLCGAK | 1486 | 1500 | 76,61 | FPIEDDKIDCLCGAK | 1486 | 1500 | 74,71 |

**Table S2.** List of selected proteins identified by mass spectrometry of HA::WDR-5.1 immunoprecipitations obtained from embryos carrying the wild type *set-2* or the *set-2(bn129)* allele, and their mammalian homologue.

| | protein name | | HA::WDR-5.1 IP, SC (number/sample) | | |
|---|---|---|---|---|---|
| | *C. elegans* | *H. sapiens* | *set-2(+)* * | *set-2(bn129)* | control |
| SET2/SET1 complex | CFP-1 | CFP1 | 18 | 0 | 0 |
| | SET-2 | SET1 | 14 | 0 | 0 |
| | SWD-2.1 | WDR82 | 2 | 0 | 0 |
| SET16/MLL complex | SET-16 | MLL1-4 | 133 | 8 | 0 |
| | PIS-1 | PTIP | 58 | 22 | 0 |
| | UTX-1 | UTX | 57 | 10 | 0 |
| core complex | RBBP-5 | RBBP5 | 42 | 15 | 0 |
| | ASH-2 | ASH2 | 29 | 6 | 0 |
| | WDR-5.1 | WDR5 | 341 | 76 | 4 |
| | DPY-30 | DPY30 | 2 | 1 | 0 |

* data previously published in Beurton et al. Nucleic Acids Res. 2019 (1)

**Table S3.** Summary of adult lifespan data presented in this work.

| Strain | Mean Lifespan±SE (days) | Maximum Lifespan | Number of worms (N) | Change (mean lifespan) | Bonferroni P-value (*vs* wt) | Figure in text |
|---|---|---|---|---|---|---|
| wt | 18.48±0.43 | 30 | 153 | | | Fig. 3A |
| *set-2(ok952)* | 15.22±0.46 | 33 | 195 | -17.60% | 0.0018 | |
| *set-2(syb2085)* | 13.68±0.51 | 32 | 167 | -26% | <0.0001 | |
| *set-2(bn129)* | 14.17±0.5 | 39 | 178 | -23% | <0.0001 | |
| *cfp-1(tm6369)* | 15.09±0.51 | 41 | 180 | -18.30% | <0.001 | |
| wt | 18.45±0.36 | 30 | 185 | | | Not shown |
| *set-2(ok952)* | 16.11±0.39 | 34 | 189 | -12.70% | 0.0007 | |
| *set-2(syb2085)* | 15.25±0.39 | 25 | 190 | -17.30% | <0.0001 | |
| *daf-2(e1370)* | 40.22±1.05 | 62 | 188 | 118% | <0.0001 | |
| wt | 19.91±0.52 | 34 | 127 | | | Not shown |
| *set-2(ok952)* | 16.96±0.6 | 35 | 136 | -14.81% | 0.0347 | |
| *set-2(bn129)* | 14.03±0.51 | 32 | 143 | -29.53% | <0.0001 | |
| *cfp-1(tm6369)* | 14.06±0.41 | 29 | 141 | -29.38% | <0.0001 | |
| wt | 15.73±0.31 | 28 | 185 | | | Not shown |
| *set-2(ok952)* | 13.9±0.34 | 23 | 175 | -11.63% | 0.0018 | |
| *set-2(syb2085)* | 12.99±0.41 | 28 | 195 | -17.40% | <0.0001 | |
| *cfp-1(tm6369)* | 13.81±0.34 | 29 | 173 | -12.20% | <0.001 | |
